## Supplementary Material for "ARCTIC-3D: Automatic Retrieval and ClusTering of Interfaces in Complexes from 3D structural information"

Marco Giulini, Rodrigo V. Honorato, Jesús L. Rivera, Alexandre M.J.J. Bonvin

### 1 Applying arctic3d-resclust to CPORT predictions

As an example scenario for the `arctic3d_resclust` CLI, we retrieve a set of possibly interacting residues for pdb file 3HMR (1) using CPORT (2) with default settings. CPORT is a software dedicated to the prediction of protein-protein interface amino acids by combining up to six different predictors. Upon providing `arctic3d_resclust` with the structure and the list of possibly interacting amino acids, we obtain two clusters (see Table 1 and Fig. 1) containing two spatially separated sets of residues.

| Group | Residues |
|---|---|
| Full set | 39,40,41,42,44,45,57,58,59,60,61,62,76,77,78,79,<br>80,82,83,84,86,98,101,105,123,124,125,126,127 |
| Cluster 1 | 39,40,41,42,44,45,105,123,124,125,126,127 |
| Cluster 2 | 57,58,59,60,61,62,76,77,78,79,80,82,83,84,86,98,101 |

**Supplementary Table 1:** Example application of `arctic3d_resclust` to the PDB structure 3HMR. The set of residues provided by CPORT is divided in two groups using average linkage hierarchical clustering (3) with a threshold of $15$ Å.

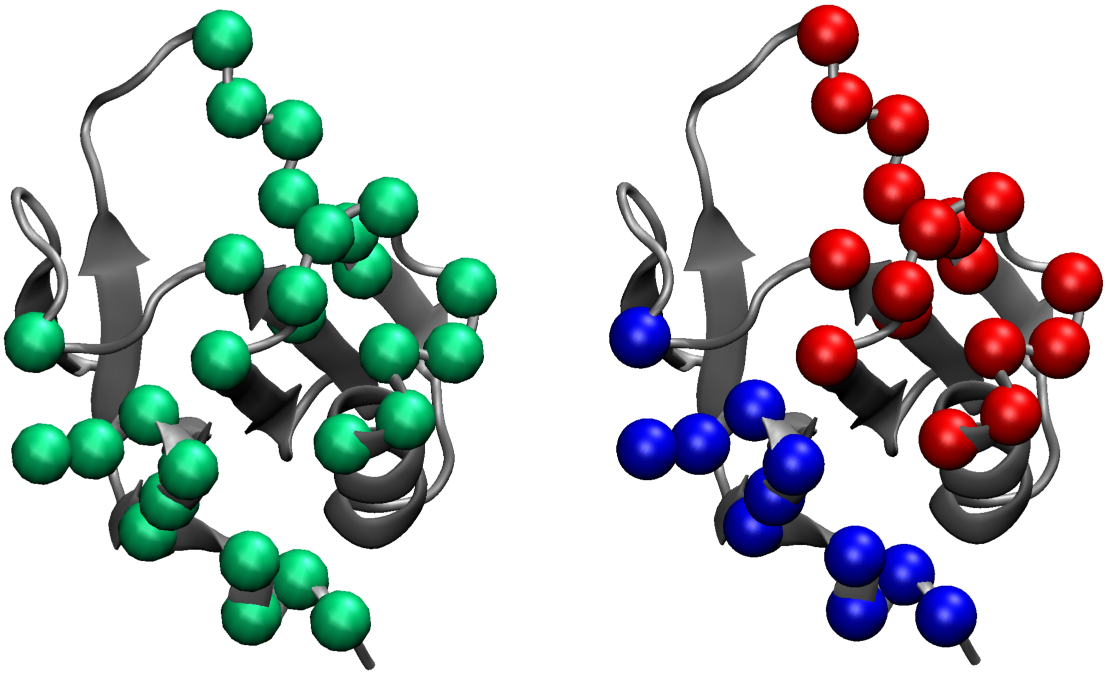

**Supplementary Figure 1:** Pictorial representation of the residue-based clustering performed by `arctic3d_resclust`. The amino acids labelled as likely interacting by CPORT are subdivided in two clearly separated patches on the protein surface. Image produced with VMD (4).

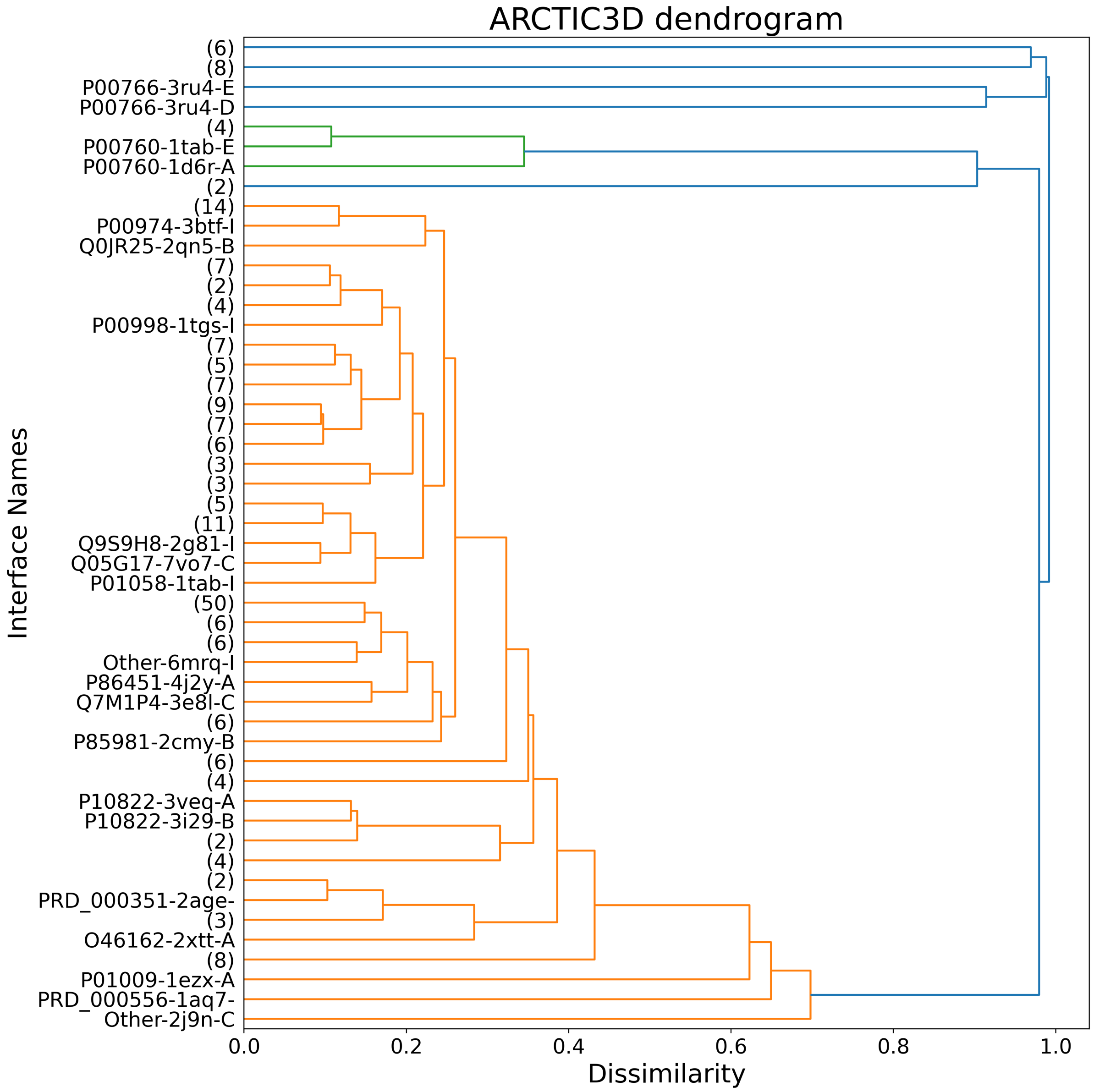


**Supplementary Figure 2:** Example dendrogram produced by ARCTIC-3D over the $228$ retained interfaces of P00760. On the x-axis we can observe how the values of dissimilarity approach $1$ as the inter-cluster distance approaches the maximum possibile dissimilarity. On the y-axis we can see some of the interface names (assigned with the `partner-pdb-chain` logic). Labels with round brackets (for example $(6)$) refer to group of interfaces that are already clustered together at that level of dissimilarity (typically very similar interfaces coming from the same PDB file).
